# Symmetric brain-liver circuits mediate lateralized regulation of hepatic glucose output in mice

**DOI:** 10.1101/2025.10.27.684732

**Authors:** Zhonglong Wang, Xiangfei Gong, Li Jiang, Ke Wang, Xinyuan Sun, Yingxi Li, Mengtong Ran, Yanshen Chen, Hongdong Wang, Xuehui Chu, Shun Wang, Junjie Wang, Xiao Zheng, Haiping Hao, Hao Xie

**Affiliations:** College of Pharmacy, China Pharmaceutical University, Nanjing 210009, China; State Key Laboratory of Natural Medicines, Institute of innovative Drug Discovery and Development, Jiangsu Provincial Key Laboratory of Targetome and Innovative Drugs, China Pharmaceutical University, Nanjing, 210009, China; Department of Endocrinology, Endocrine and Metabolic Disease Medical Center, Nanjing Drum Tower Hospital, Affiliated Hospital of Medical School, Nanjing University, Nanjing, 210008, China; Department of Pancreatic and Metabolic Surgery, Nanjing Drum Tower Hospital, Affiliated Hospital of Medical School, Nanjing University, Nanjing, 210008, China

**Keywords:** symmetric regulation, intra-liver sympathetic innervation, hepatic glucose production

## Abstract

Neural lateralization is well recognized in the control of contralateral somatic movement, yet its relevance to visceral organ regulation remains poorly understood. This study aimed to determine whether the central nervous system exerts lateralized control over hepatic glucose metabolism and to localize the site of peripheral sympathetic crossover to the liver. Pseudorabies virus (PRV) tracing revealed bilateral projections from the lateral paragigantocellular nucleus (LPGi) with preferential innervation of the contralateral hepatic lobes. Unilateral LPGi activation elevated systemic glucose by enhancing glycogenolysis and gluconeogenesis specifically in contralateral lobes, whereas bilateral activation produced additive effects. Following unilateral hepatic denervation, contralateral LPGi activation induced metabolic compensation in the remaining innervated lobes, characterized by increased norepinephrine release, glucose production, and glycogen depletion. Whole-mount tissue clearing and dual viral tracing localized the sympathetic crossover to the porta hepatis. Developmental analysis showed that lobar-specific innervation along the vasculature emerges by postnatal week 2. These findings demonstrate that the brainstem can exert lobe-specific, lateralized control of hepatic glucose metabolism via bilaterally projecting brain – liver sympathetic pathways. This contralateral regulation arises from a peripheral decussation at the porta hepatis, and the compensatory activation observed after denervation reveals an intrinsic neuroadaptive mechanism that helps safeguard systemic glucose homeostasis.

**Highlights:**

- Brain-liver sympathetic projections exhibit predominant contralateral innervation
- Unilateral LPGi activation drives glucose production in contralateral hepatic lobes
- Unilateral denervation augments contralateral LPGi-mediated metabolic compensation
- Sympathetic crossover to the liver localizes at the porta hepatis

## Introduction

Pronounced functional lateralization within the central nervous system (CNS) is a well-documented phenomenon,^(1,2)^ traditionally associated with cognitive and motor processes such as language, voluntary movement, and spatial navigation.^(3,4)^ Emerging evidence indicates that this lateralization extends beyond cortical functions to the autonomic regulation of visceral organs, thereby enhancing the efficiency of physiological responses.^(5–7)^ Such CNS specialization allows for optimized resource allocation and context-dependent prioritization of functional outputs. For instance, paired organs such as kidneys and adrenal glands receive predominantly contralateral neural inputs, enabling coordinated bilateral function.^(8,9)^ Previous studies have shown that the left hemisphere preferentially modulates sympathetic outflow to the right kidney and *vice versa*,^(8,10)^ and that cortical projections to the adrenal medulla are primarily contralateral, with the rostral cingulate motor area as a bilateral exception.^(9)^ However, the influence of neural lateralization on structurally asymmetric visceral organs remains incompletely understood and appears to be organ-specific. The heart, for instance, exhibits distinct lateralized autonomic innervation: left-sided fibers predominantly mediate chronotropic regulation via the sinoatrial node, whereas right-sided fibers exert greater inotropic and dromotropic functions.^(6,11)^ Additionally, cortical neurons regulating gastric sympathetic function are predominantly localized in the right hemisphere, whereas parasympathetic control displays only a modest right-sided predominance.^(7,12)^ These examples underscore the organ-specific nature of lateralized autonomic neural projections.

The liver, as the largest asymmetrical visceral organ, plays a central role in systemic metabolic homeostasis,^(13,14)^ orchestrating physiological processes such as glucose production, lipid metabolism, protein synthesis, and detoxification.^(16,17)^ In mice, hepatic innervation first appears in extrahepatic bile ducts at embryonic day 17.5 and extends into lobes during the early postnatal period,^(18)^ yet the precise pattern of this innervation remains unknown. Previous anatomical and functional studies, including our own, have established that the liver receives dense sympathetic innervation from postganglionic neurons of the celiac-superior mesenteric ganglia (CG-SMG).^(19,20,21,22)^ Within this context, the brainstem lateral paragigantocellular nucleus (LPGi) has been identified as a key node in the regulation of hepatic sympathetic tone and glucose output.^(20)^ However, whether this regulation is lateralized and lobe-specific, and where sympathetic crossover occurs, has not been systematically defined.

To address these gaps, we applied retrograde trans-synaptic tracing with pseudorabies virus (PRV) to delineate LPGi projections to hepatic lobes and assess their lateralization.^(21–23)^ Functional experiments employing chemogenetic and optogenetic activation of LPGi neurons clarified the physiological impact of this neural circuitry on hepatic glucose production (HGP). Additionally, we identified compensatory neural mechanisms that maintain systemic glucose homeostasis following unilateral disruption of the neural circuit regulating hepatic innervation. Whole-mount clearing identified the porta hepatis as the site of sympathetic crossover and revealed developmental axon extension of hepatic fibers along the vasculature. Together, these findings reveal that lateralized neuronal circuits exert lobe-specific control over hepatic glucose output and recruit neuroadaptive responses to preserve metabolic homeostasis following unilateral disruption.

## Results

### 1. Contralateral projections from the LPGi to distinct hepatic lobes

Previous studies of intrahepatic innervation mainly focused on individual lobes, such as the left or right lateral lobes in mice.^(14,15)^ We first asked whether sympathetic innervation is a uniform feature across all lobes. To address this, we examined the intact mouse liver using tissue clearing combined with immunofluorescence staining of tyrosine hydroxylase (TH), the rate-limiting enzyme in catecholamine synthesis. This analysis revealed that, although hepatic lobes differ in volume (Figure 1A), sympathetic fibers consistently branch into all lobes with comparable density (Figure 1A). To further investigate the anatomical basis of central neural regulation of the liver, we performed multi-synaptic retrograde tracing by injecting PRV into individual hepatic lobes, including the left lateral, median, right posterior, right anterior, and caudate lobes (Figures S1A and S1B). Five days post-injection, PRV-labeled neurons were consistently identified in the LPGi of the brainstem (Figure 1B, Figures S1C-S1F). Quantitative analyses revealed a clear contralateral dominance: the injection into left lateral and median lobes yielded approximately twice as many labeled neurons in the right LPGi as in the left, whereas the injection into right posterior, right anterior, and caudate lobes preferentially labeled neurons in the left LPGi (Figure 1C). These findings provide direct anatomical evidence of a symmetric projection pattern in which each LPGi hemisphere preferentially innervates contralateral hepatic lobes (Figure 1D). Based on their robust lateralization, the right anterior and median lobes were selected as representative targets for subsequent analyses.

**Figure 1.**
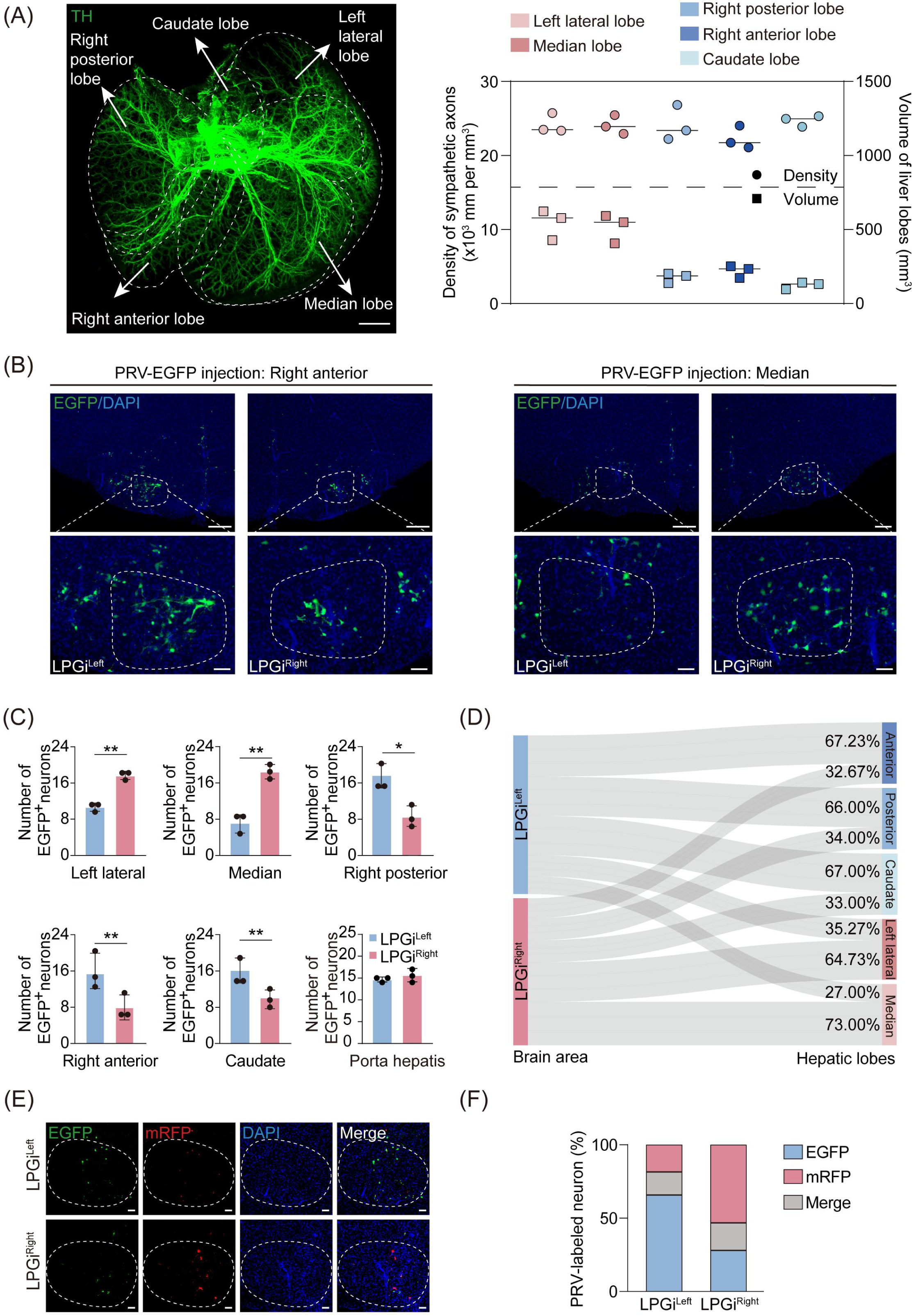
The brain contralaterally projects to the liver. (A) Representative 3D images of TH-immunostained mouse liver (left), with quantification of sympathetic axon density and lobe volume (right). Scale bar, 3,000 μm. (B) Representative immunofluorescence images of PRV-labeled neurons (EGFP) in the left and right LPGi following PRV injections into the right anterior (left) and median (right) lobes. Scale bars, 200μm (top) and 100 μm (bottom). (C) Quantification of PRV-labeled neurons in left and right LPGi across different hepatic lobes: left lateral, median, right posterior, right anterior, caudate, and porta hepatis (*n* = 3). (D) Sankey diagram showing projection patterns from left and right LPGi to individual hepatic lobes. (E and F) Representative slices of EGFP^+^ and mRFP^+^ neurons in left (top) and right (bottom) LPGi following PRV-EGFP (right anterior lobe) and PRV-mRFP (median lobe) injections. Proportions of EGFP^+^, mRFP^+^, and co-labeled neurons in left and right LPGi (F, *n* = 3). Scale bars, 100 μm. Data are expressed as means ± SEM, with individual values shown for all experiments. \**p* < 0.05, \*\**p* < 0.01. Unpaired Student’s t-test for (C).

To validate these projection patterns, we performed dual labeling by co-injecting PRV-CAG-EGFP into the right anterior lobe and PRV-CAG-mRFP into the median lobe. In the left LPGi, 73.7% of PRV-positive neurons expressed EGFP, showing projections to the right anterior lobe, whereas in the right LPGi, 62.5% expressed mRFP, indicating projections to the median lobe (Figures 1E and 1F). A subset of neurons displayed co-labeling, suggesting that although the predominant projection pattern is contralateral, a population of LPGi neurons innervates both lobes bilaterally. These findings establish a contralaterally biased projection pattern from each LPGi to distinct hepatic lobes, providing the anatomical foundation for lateralized brain-liver circuits that regulate hepatic glucose metabolism.

Because PRV tracing cannot distinguish sympathetic efferent from vagal sensory pathways, we next sought to determine the circuit identity of liver-projecting LPGi neurons. Unlike the NTS, a well-established hepatic sensory center served here as a positive control,^(24)^ the LPGi contained few CGRP-positive cell bodies (Figure S1G), indicating a lack of peptidergic sensory projections. Moreover, co-injection of hSyn-cre and DIO-Axon-EGFP into the LPGi did not yield Axon-EGFP-positive signals in either the dorsal root ganglia (DRG) or nodose ganglia (NG) (Figures S1H-S1J). Together, these findings indicate that the LPGi specifically regulates sympathetic, rather than vagal sensory, inputs to the liver.

### 2. Differential HGP by LPGi lateralization

To assess the functional implications of this anatomical laterality, we examined whether activation of the LPGi in each hemisphere modulates HGP in a lobe-specific manner. Prior single-nucleus RNA sequencing and immunofluorescence analyses demonstrated that liver-projecting LPGi neurons are predominantly GABAergic,^(45)^ we selectively activated these neurons unilaterally or bilaterally using chemogenetics (Figure 2A, Figure S2A). LPGi activation significantly increased blood glucose, with area under the curve (AUC) increases by 27%, 34%, and 52% following left-, right-, and bilateral stimulation, respectively (Figure 2A). Moreover, right-sided activation induced greater hyperglycemia than left-sided stimulation, possibly reflecting differences in the metabolic capacity or volume of the contralateral hepatic lobes (Figure 1A).

**Figure 2.**
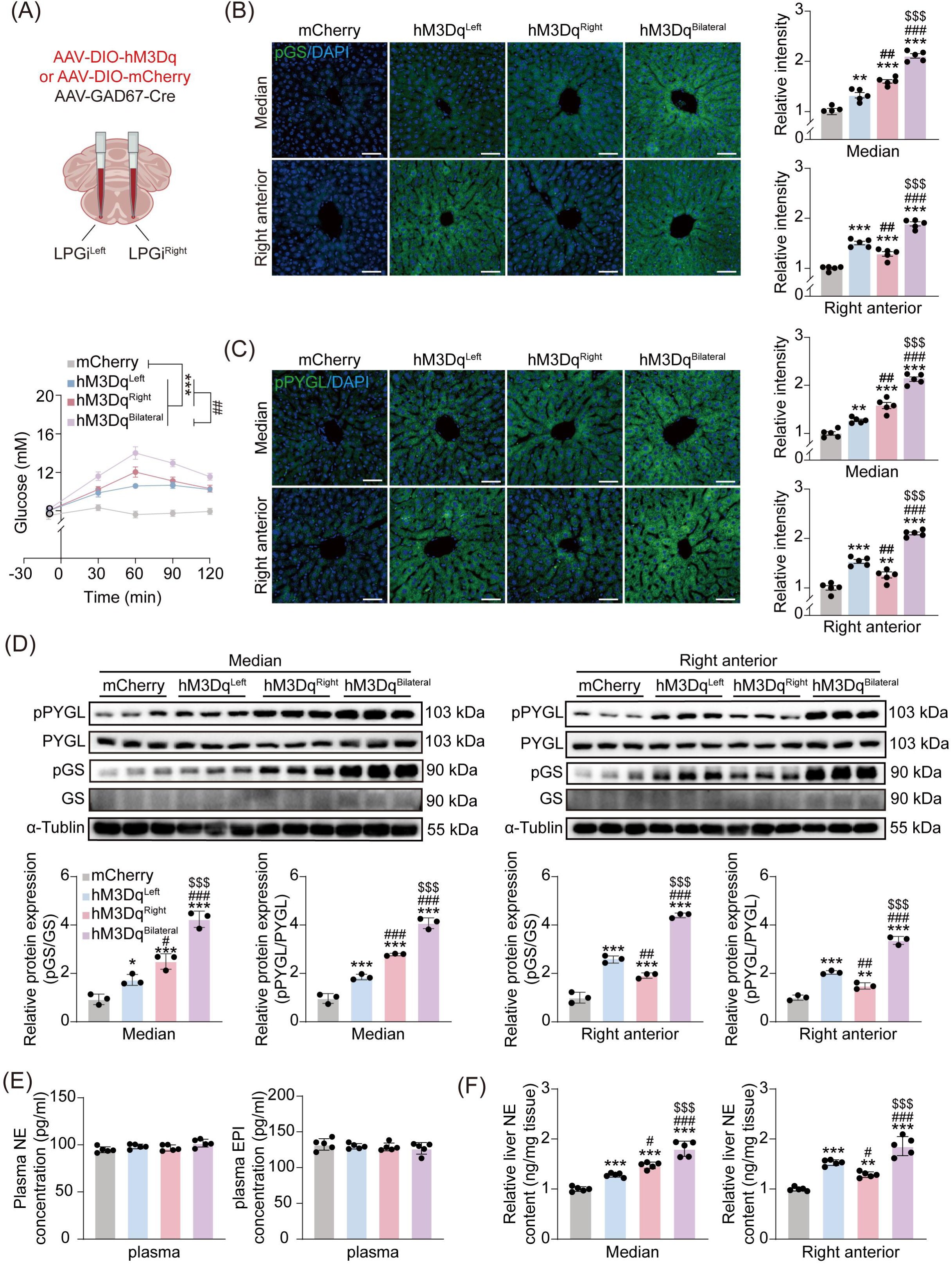
LPGi stimulation raises blood glucose by driving contralateral, lobe-specific hepatic glycogenolysis. (A) Scheme for injecting AAV-DIO-hM3Dq or AAV-DIO-mCherry (top). Blood glucose levels following chemogenetic activation of the left, right, or bilateral LPGi, measured at −10, 30, 60, 90, and 120 min (bottom, *n* = 6). (B and C) Representative immunofluorescence images of pGS (B) and pPYGL (C) expression in the median (top) and right anterior (bottom) lobes after 1 h chemogenetic activation of left, right, or bilateral LPGi, with quantification of relative fluorescence intensity (*n* = 4-5). Scale bars, 100 μm. (D) Western blot analysis of pPYGL, PYGL, pGS, and GS in median (left) and right anterior (right) lobes after 1 h activation, with densitometric quantification (bottom, *n* = 3). (E) Plasma concentrations of norepinephrine (NE) (left, *n* = 5) and epinephrine (EPI) (right, *n* = 5) following 1 h activation. (F) NE contents in median (left, *n* = 5) and right anterior (right, *n* = 5) lobes after 1 h activation. Data are expressed as means ± SEM, with individual values shown for all experiments. \**p* < 0.05, \*\**p* < 0.01, \*\*\**p* < 0.001; #*p* < 0.05, ##*p* < 0.01, ###*p* < 0.001; $$$*p* < 0.001. One-way ANOVA with the Tukey test for (A -F).

At the molecular level, one hour following stimulation, phosphorylation of glycogen phosphorylase (pPYGL) and glycogen synthase (pGS) increased in contralateral lobes, as assessed by immunofluorescence and western blot. Left LPGi activation preferentially enhanced pPYGL and pGS levels in the right anterior lobe compared with right-sided activation, whereas right LPGi activation exerted a greater effect in the median lobe relative to left-sided stimulation (Figures 2B-2D). Bilateral stimulation further amplified pPYGL and pGS expression in both lobes compared with unilateral activation. PAS staining confirmed glycogen depletion in metabolically engaged contralateral lobes (Figure S2B). In terms of gluconeogenesis, enzymatic activity assays revealed increased phosphoenolpyruvate carboxykinase (PEPCK) and glucose-6-phosphatase (G6PC) activity in hepatic lobes undergoing enhanced glycogenolysis, despite unchanged protein levels within the one-hour window (Figures S2C-S2F).

To exclude contributions from systemic endocrine responses, particularly those mediated by the hypothalamic-pituitary-adrenal (HPA) axis, we measured plasma NE, EPI, insulin, and glucagon 1 hour after LPGi activation. None of these hormones was significantly altered by either unilateral or bilateral stimulation (Figure 2E and Figure S2G). In contrast, intrahepatic NE content rose specifically in contralateral lobes: by 1.49-fold in the right anterior lobe after left LPGi activation, 1.53-fold in the median lobe after right LPGi activation, and over 1.85-fold in both lobes after bilateral activation (Figure 2F). These findings support a mechanism in which lobe-specific sympathetic activation drives hepatic glucose output in a lateralized manner, independent of systemic hormonal changes.

To achieve higher temporal precision, we applied optogenetics by delivering AAV9-GAD67-Cre with AAV9-DIO-ChR2-mCherry viruses to the LPGi (Figure S3A, Figure S4A). Compared with chemogenetics, optogenetic activation elicited greater hyperglycemia, with AUC rising by 32%, 42%, and 62% for left-, right-, and bilateral stimulation, respectively (Figure S3A). Molecular and histological analyses paralleled those from chemogenetics, including contralateral and additive effects on pPYGL and pGS (Figures S3B-S3D), glycogen depletion (Figure S4B), elevated intrahepatic NE content (Figure S3F), and enhanced PEPCK and G6PC activity (Figures S4C-S4F), without changes in plasma catecholamines (Figure S3E). Taken together, these findings demonstrate that each LPGi hemisphere preferentially drives glucose production in contralateral hepatic lobes, primarily through localized sympathetic activation and glycogenolysis in contralateral hepatic regions.

### 3. Compensatory neural responses following unilateral hepatic denervation

While compensatory adaptations following unilateral damage are well described in symmetrical organs, it remains unclear whether similar mechanisms operate in asymmetrical organs such as the liver.^(26–28)^ To test this, we chemically denervated the right- or left-sided lobes using 6-OHDA. Despite the absence of directly sympathetic input to denervated lobes, systemic blood glucose levels were unchanged compared with controls (Figures 3A-3B, Figure S5A), indicating functional compensation through the remaining intact liver. Further analyses revealed increased expression of pPYGL and pGS, as well as elevated intrahepatic NE levels and glycogen depletion in non-denervated hepatic lobes (Figures 3C-3H, Figures S5B-S5E).

**Figure3.**
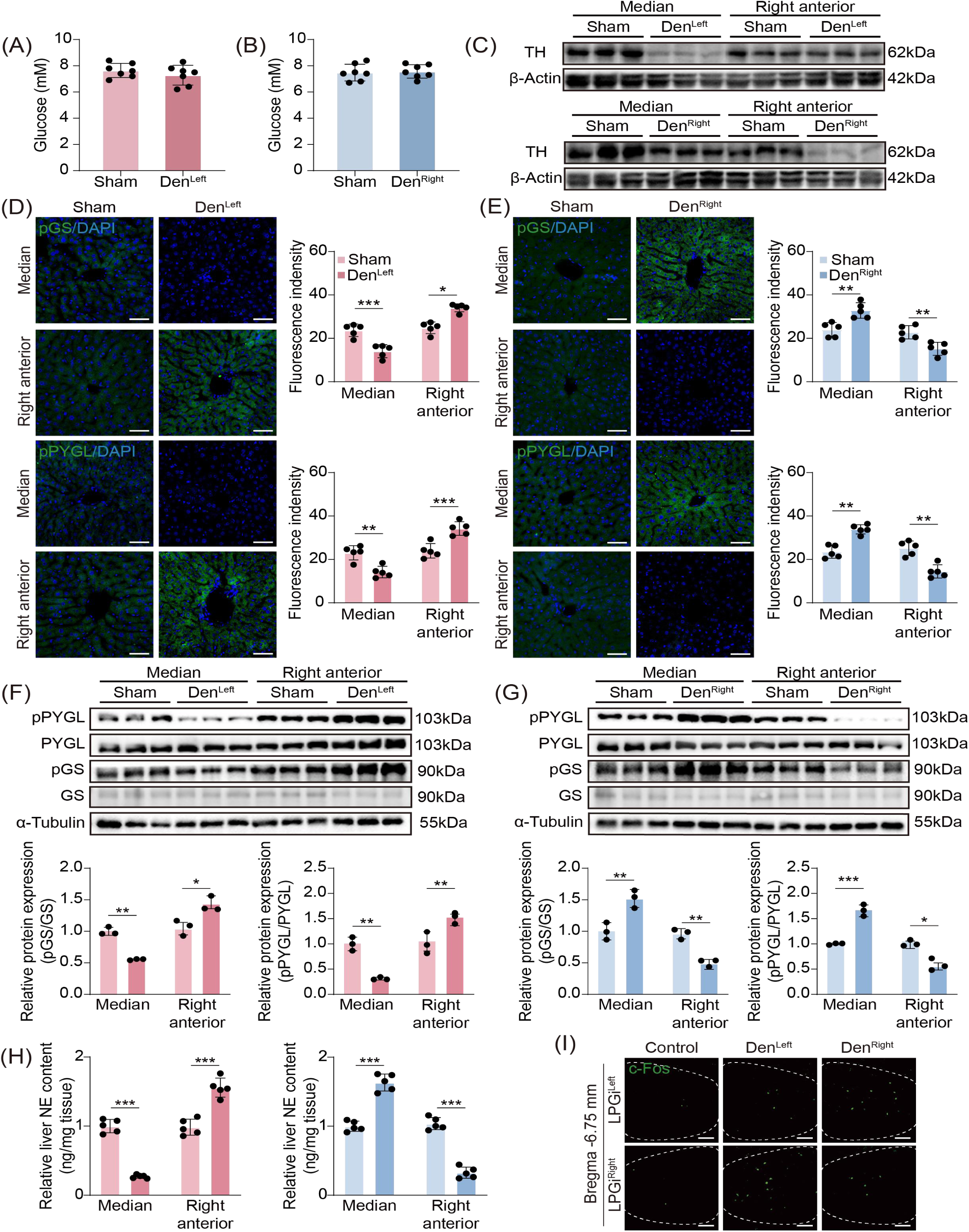
LPGi maintains glucose balance through compensatory use of ipsilateral hepatic lobes under contralateral lobe impairment. (A and B) Blood glucose levels in the left- (A, n = 7) or right-sided (B, n = 7) lobes denervated mice. (C) Western blot of TH protein in median and right anterior lobes after denervating left- (top) or right-sided (bottom) lobes (n = 3). (D and E) Representative immunofluorescence images of pGS (top) and pPYGL (bottom) expression in right anterior and median lobes in left- (D) or right-sided (E) lobes denervated mice, with quantification of relative fluorescence intensity (right, n = 5). Scale bars, 100 μm. (F and G) Western blot analysis of pPYGL, PYGL, pGS, and GS proteins in median and right anterior lobes (top) in left- or right-sided lobes denervated mice, with densitometric quantification (bottom, n = 3). (H) NE contents of median and right anterior lobes in left- or right-sided lobes denervated mice (n = 6). (I) Representative immunofluorescence images of c-FOS expression in left and right LPGi in left- or right-sided lobes denervated mice. Scale bars, 100 μm. Data are presented as means ± SEM, with individual values shown for all experiments. *p < 0.05, **p < 0.01, ***p < 0.001. Unpaired Student’s t-test for (A-B, D-H)

To elucidate the underlying mechanism, we quantified c-FOS expression in the LPGi following unilateral hepatic denervation. Right-sided lobes denervation increased c-FOS in the left LPGi, and *vice versa* (Figure 3I), suggesting that contralateral LPGi upregulates its sympathetic output to intact ipsilateral hepatic lobes, thereby compensating for the loss of innervation in denervated lobes. Together, these findings identify a neuroadaptive mechanism in which the ipsilateral LPGi augments sympathetic output to intact hepatic lobes, thereby maintaining systemic glucose homeostasis despite partial intrahepatic neural disruption.

### 4. Peripheral decussation of sympathetic hepatic control at the porta hepatis

Contralateral neural circuitry typically arises from central decussations, such as corticospinal tracts that cross at the pyramids for limb motor control,^(29,30)^ and visual pathways that decussate at the optic chiasm for hemispheric processing of visual fields.^(31)^ Whether the brain’s contralateral regulation of the liver occurs centrally or peripherally, however, has remained unknown. Using whole-mount clearing, we visualized the brain–liver sympathetic circuit and found that preganglionic neurons in the thoracic spinal cord (T6 – T12) send descending fibers via the splanchnic nerves to innervate postganglionic neurons in the CG-SMG (Figure 4A). Further whole-mount TH immunostaining showed that TH-positive sympathetic cell bodies within the CG-SMG project to the liver along the hepatic vasculature (Figure 4B and Figure S5F). These observations suggest that the nerve bundles likely decussate at the porta hepatis before entering the individual hepatic lobes.

**Figure 4.**
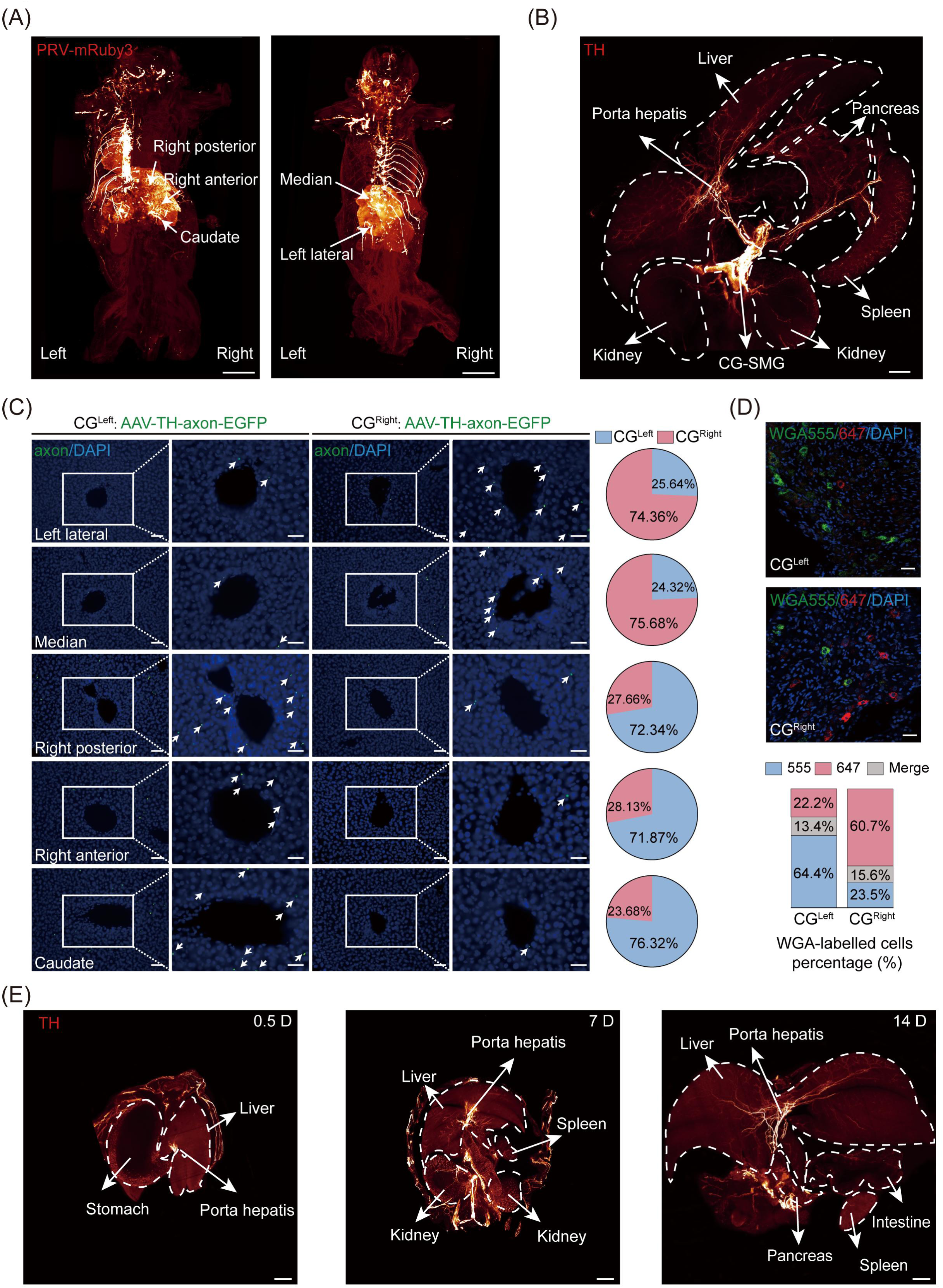
Sympathetic projections to the liver decussate at the porta hepatis, progressively formed during development. (A) Representative 3D images of mice following PRV injection into right- (left) or left-sided (right) lobes. Scale bars, 3000 µm. (B) Representative 3D images of CG-SMG projections to peripheral organs. Scale bar, 3,000 μm. (C) Representative images of sympathetic axons after unilateral CG injection, with quantification of EGFP-labeled axon distribution across hepatic lobes. Scale bars, 100 μm. (D) Representative images of WGA-labeled neurons in left and right CG, with percentages of WGA555^+^, WGA647^+^, and co-labeled neurons. Scale bars, 100 μm. (E) Representative images showing the developmental progression of CG-SMG projections to the liver from postnatal week 0 to week 2. Scale bars, 3,000 µm.

To test this, we performed unilateral anterograde tracing by injecting AAV-TH-DIO-EGFP into a single CG (Figure S5G). This revealed a strictly contralateral innervation pattern: the left CG mainly projected to the right posterior, right anterior, and caudate lobes, whereas the right CG prominently innervated the left lateral and median lobes (Figure 4C). To validate this pattern, we performed dual retrograde tracing by injecting Alexa Fluor 647- and 555-conjugated wheat germ agglutinin (WGA) into left- and right-sided lobes, respectively (Figure S5H). Consistent with the anterograde results, left CG neurons predominantly projected to the right-sided lobes, while right CG neurons targeted the left-sided lobes, confirming robust contralateral preference (Figure 4D). Thus, these findings demonstrate that central inputs descend via the SyC to the CG-SMG complex, where nerve bundles converge and extend along the portal vein, branching within the porta hepatis to innervate individual hepatic lobes.

To explore the development of the intrahepatic neural network in mice, we therefore performed tissue clearing on the torsos of mice from postnatal weeks 0 to 2. We observed that at week 0, the sympathetic nerve bundles were located at the porta hepatis but had not yet extended into the liver (Figure 4E, left). By week 1, the sympathetic nerves had begun to grow into the liver (Figure 4E, middle). It was not until week 2 that the sympathetic nerves fully infiltrated all liver lobes (Figure 4E, right). These findings indicate that, after the first postnatal week, sympathetic nerves grow from the porta hepatis into the liver lobes and form a well-developed sympathetic neural network by the second week.

## Discussion

The CNS exhibits functional asymmetry despite structural bilateral symmetry, a phenomenon well recognized in language, handedness, and spatial cognition.^(32,33)^ Recent advances extend this concept beyond cortical domains to central autonomic regulation of peripheral organs.^(34,35)^ Trans-neuronal tracing of symmetrical organs such as the kidneys and thymus has revealed contralateral control with bilateral coordination,^(8,35)^ whereas the lateralized regulation of asymmetrical organs like the liver, heart, and stomach remains poorly defined and conflicting.^(6,12,37)^ For instance, parasympathetic cardiac control has been reported to exhibit left-hemispheric dominance,^(6)^ yet clinical evidence revealed that arrhythmias may arise after strokes in either hemisphere.^(38)^ These discrepancies highlight the complexity of lateralized autonomic control and the intrinsic bilaterality within the autonomic network. In this study, we systematically mapped and functionally interrogated the laterality of brain-liver circuitry, focusing on the LPGi. Our anatomical and functional experiments provide compelling evidence that although a subset of LPGi neurons project bilaterally, the predominant organization is contralateral and lobe-specific.

Recent advances in viral tracing and whole-tissue imaging have greatly accelerated our understanding of peripheral neuroanatomy.^(39–41)^ For instance, distinct gastrointestinal regions are differentially regulated by subdivisions within the CG-SMG,^(19)^ while the kidneys predominantly receive ipsilateral input from sympathetic postganglionic neurons residing in the renal sympathetic ganglia.^(20)^ In contrast, components of the immune system, such as the thymus, exhibit lateralized central regulation, with each hemisphere preferentially modulating the contralateral thymic lobe.^(36)^ Using PRV-mediated retrograde tracing with whole-mount clearing, we mapped a brain-liver axis arising from the LPGi and revealed a strikingly lateralized projection pattern: left LPGi neurons preferentially innervated right-sided lobes, whereas right LPGi neurons predominantly targeted left-sided lobes. A small subset of LPGi neurons was double-labeled after bilateral PRV injections, suggesting a fraction of these neurons projects bilaterally to both sides of the liver. Such neurons may support interlobar coordination. Together, this high-resolution anatomical mapping provides direct evidence for lateralized central projections to an asymmetric organ and reveals a previously unrecognized dimension of laterality in autonomic liver innervation.

Functionally, this anatomical organization translates into lobe-specific regulation of HGP. Sympathetic innervation of the liver is well established as a major driver of glucose output, with prior electrical stimulation studies demonstrating robust hepatic glucose release in response to sympathetic activation.^(42–44)^ Consistent with these observations, both chemogenetic and optogenetic activation of LPGi GABAergic neurons significantly elevated systemic glucose levels by promoting glycogenolysis and gluconeogenesis in the contralateral lobes. These metabolic responses were accompanied by localized NE release, increased phosphorylation of key metabolic enzymes (pPYGL and pGS), and glycogen depletion.^(45,46)^ Bilateral LPGi activation produced additive effects, indicating that both sides of the brainstem can cooperatively regulate hepatic metabolism in a spatially segregated manner. This pattern suggests that hepatic glucose output can be modulated in a lobe-specific, rather than uniform whole-organ, manner. This spatial organization is likely obscured by conventional electrical stimulation, which indiscriminately activates heterogeneous sympathetic fibers.^(44,45)^ By contrast, chemogenetic and optogenetic approaches permit selective, cell type-specific manipulation of LPGi neurons, thereby revealing the contralateral and lobe-specific architecture of brain-liver sympathetic control.

Beyond identifying lateralized control, we also reported a compensatory mechanism following unilateral hepatic denervation. When neural input from one side of the brainstem was attenuated, compensatory neural plasticity emerged through increased sympathetic drive from contralateral LPGi, enhancing gluconeogenesis and glycogenolysis within the remaining innervated hepatic lobes. This response was supported by increased c-FOS expression in contralateral LPGi, higher intrahepatic NE content, and elevated pPYGL and pGS expression in unaffected lobes. Such plasticity parallels compensatory phenomena in other systems, including contralateral enhancement after renal dysfunction and cortical dominance shifts in patients with unilateral hearing loss.^(26,47)^ Although enhanced sympathetic output appears to mediate much of this compensation, our findings suggest that the underlying regulation extends beyond a purely descending pathway. In particular, c-FOS activation in the contralateral LPGi after unilateral 6-OHDA-mediated denervation suggests that the loss of peripheral input may be sensed through an ascending neural pathway, centrally integrated, and translated into compensatory sympathetic output to the intact hepatic lobes. These results therefore support a model in which hepatic glucose production is regulated by an integrated afferent-central-efferent loop, with our current analyses primarily resolving its efferent component.

We further show that contralateral regulation of hepatic metabolism arises from a peripheral decussation, challenging the long-held view that such organization arises exclusively from central crossings. Unlike somatic and sensory systems, where contralateral control is mediated by pyramidal or optic decussations in CNS nuclei,^(2,28,29)^ hepatic sympathetic projections descend ipsilaterally from the brainstem to the CG-SMG before crossing to innervate contralateral lobes. This peripheral arrangement suggests that spatially organized control can be implemented downstream of peripheral ganglia, representing a potential evolutionary adaptation. More broadly, it calls for re-examination of neural laterality paradigms and identifies peripheral ganglia as decision nodes and therapeutic targets for organ-specific neuromodulation.

Despite these advances, several limitations warrant mention. First, the neuronal subtypes within the LPGi that mediate contralateral projections remain unidentified. Future research combining genetic labeling with single-cell RNA sequencing will be needed to resolve their molecular and functional identities. Second, technical constraints prevented simultaneous bilateral viral tracing and whole-mount mouse imaging in the same animal, restricting direct assessment of interhemispheric interactions. Third, although whole-mount TH immunostaining with three-dimensional reconstruction revealed sympathetic nerve bundles projecting from the CG to the liver, TH is not entirely specific and can also label a subset of sensory neurons. More selective approaches, such as genetic targeting of sympathetic lineages, will be important for further validation. Fourth, although our data support a peripheral decussation at the porta hepatis, direct validation of this crossover site was not feasible with local pharmacological blockade, as currently available approaches lack sufficient spatial specificity and would likely perturb multiple neural components. Future studies employing more selective inhibitory strategies will be required to directly test this possibility. Finally, although our murine data provide new insights into lateralized autonomic control of hepatic metabolism, their translational relevance to humans remains to be established. Progress in noninvasive imaging and neural circuit mapping may ultimately enable analogous studies in clinical settings.

In conclusion, this study establishes that the brain exerts lateralized and lobe-specific control over hepatic glucose metabolism, with contralateral regulation arising from a peripheral decussation at the porta hepatis, and a compensatory mechanism maintaining glucose homeostasis after unilateral hepatic injury. Developmentally, sympathetic projections began extending into the liver during the first postnatal week, forming a mature network by week 2 that enables both baseline regulation and adaptive responses. Together, these findings extend the concept of lateralization beyond cognition, motor control, and symmetric organs, with potential implications for targeted neuromodulation strategies in organ-specific diseases.

## Methods

### Animals

All animal experiments were performed in accordance with the Guide for the Care and Use of Laboratory Animals and were approved by the Institutional Animal Care and Use Committee of China Pharmaceutical University. Adult C57BL/6J mice (male, 8-week-old) were used in this study. For retrograde trans-synaptic PRV tracing, 8-week-old mice received intrahepatic viral injections and were sacrificed 5 days post-injection. For chemogenetic and optogenetic manipulations, stereotaxic AAV injections were performed at 8 weeks of age. Mice were allowed 4 weeks for viral expression and recovery before undergoing metabolic tests or light stimulation at 12 weeks of age. For chemical denervation (6-OHDA), 8-week-old mice were injected into targeted lobes and examined 7 days post-denervation. For postnatal innervation mapping, neonatal mice at postnatal week 0 (P0), week 1 (P7), and week 2 (P14) were harvested for tissue clearing. Mice were group-housed (3-5 per cage) under a standard 12-hour light/dark cycle (lights on at 7:00 AM) at a controlled temperature (22 ± 1 °C) and humidity (50 ± 10%), with ad libitum access to a standard laboratory chow diet and water.

For all surgical and terminal procedures, mice were anesthetized with isoflurane (induction 3-4%, maintenance 1.5-2% in 100% oxygen). Littermates of the same sex were randomly assigned to experimental or control groups using a random number generator. Experimenters were blinded to group allocation during data collection and analysis where feasible. All efforts were made to minimize animal suffering and to use the minimum number of animals required for statistical reliability.

### Viral vectors

For retrograde tracing of LPGi projection to individual hepatic lobes, we injected PRV-CAG-EGFP (BrainVTA, Cat#P03001) and PRV-CAG-mRFP (BrainVTA, Cat#P03002) into distinct hepatic lobes, respectively. For chemogenetic manipulation, we used a combination of AAV9-GAD67-Cre (BrainVTA, Cat#PT-6600) with AAV9-DIO-hM3Dq (BrainVTA, Cat#PT-0019) or AAV9-DIO-mCherry (BrainVTA, Cat#PT-0115). For optogenetic manipulation, we used a combination of AAV9-GAD67-Cre with AAV9-DIO-ChR2-mCherry (BrainVTA, Cat#PT-0002) or AAV9-DIO-mCherry. For anterograde tracing from the CG-SMG to hepatic lobes, we injected AAV9-TH-Cre (BrainVTA, Cat#PT-2947) with AAV9-EF1α-DIO-Axon-EGFP (BrainVTA, Cat#BC-0197) into the unilateral CG-SMG. All viral vectors were used at titers >5 × 10¹² vg/mL.

### PRV injection

For retrograde tracing from the liver, pseudorabies virus (PRV-CAG-EGFP, 5.00 ×10^9^ PFU/ml, BrainVTA, Cat#P03001) was used to identify upstream autonomic inputs to the hepatic lobes. Following laparotomy and exposure of the liver as described above, a series of focal injections was performed to restrict viral labeling to circuit-specific connections. A total volume of 1 µL of PRV per lobe was slowly pressure-injected into the targeted lobe using a glass micropipette attached to a microsyringe. Injections were made at multiple sites (2-3 sites) within each lobe, with careful coordination to avoid leakage into the peritoneal cavity or surrounding tissues. Following each injection, the micropipette was left in place for 5 min to minimize backflow, and the injection site was briefly swabbed with sterile saline. The intestines were then repositioned, and the abdominal wall was sutured. Animals were perfused 5 days post-injection, and brain sections were processed for PRV immunodetection to map infected neuronal populations.

### Quantification of PRV-labeled neurons

For quantitative analysis, serial coronal sections (40 µm thickness) encompassing the entire rostrocaudal extent of the LPGi were collected. Every third section was selected for immunostaining and imaging. Images were captured using a fluorescence microscope (Leica) with a 20×objective. PRV-positive neurons in each selected section were counted manually using the Cell Counter plugin in ImageJ software. Neurons were identified based on fluorescent signal intensity above background and characteristic neuronal morphology with clearly labeled nuclei and cytoplasm.

### Brain stereotactic injection

Mice were anesthetized with isoflurane (RWD Life Science Co., Ltd., Cat#68811). For brain injection, the coordinates of LPGi were determined using the Allen Mouse Brain Atlas (AP −6.75 mm, ML ±0.95 mm, DV −5.85 mm). A total of 200 nL of viral vector was bilaterally delivered at a rate of 50 nL/min.

### Intrahepatic sympathetic nerve tracing

Following laparotomy, the intestines were gently exteriorized and covered with saline-moistened gauze to expose the celiac ganglion-superior mesenteric ganglion (CG-SMG). For anterograde tracing of CG-SMG projections to hepatic lobes, a viral cocktail containing AAV9-TH-Cre (BrainVTA, Cat#PT-2947) and AAV9-EF1 α -DIO-Axon-EGFP (BrainVTA, Cat#BC-0197) was injected into either the left or right CG using fine glass micropipettes. A volume of 100 nL per site was delivered at a slow infusion rate of 50 nL/min to minimize tissue damage and restrict viral spread to the targeted ganglion, and tissues were collected 3 weeks post-injection. For retrograde tracing, 500 nL of 1 mg/ml Alexa Fluor 647-conjugated wheat germ agglutinin (WGA 647, Thermo, Cat#W32466) and Alexa Fluor 555-conjugated wheat germ agglutinin (WGA 555, Thermo, Cat#W32464) were injected into the left- and right-sided lobes, respectively, using an infusion rate of 100 nL/min. Tissues were harvested 3 days thereafter.

### Intrahepatic sympathetic denervation

To achieve lobe-specific sympathetic denervation, individual hepatic lobes were physically separated and isolated during the injection procedure to minimize diffusion of the neurotoxin to adjacent tissue. The injection volume was calculated based on the proportion of each lobe relative to the total liver weight. A solution of 6-hydroxydopamine (6-OHDA, 12 mg/ml in sterile saline containing 0.01% ascorbic acid; Sigma, Cat#162957) was injected into each target lobe at a volume of 2,500-3500 nL and a flow rate of 300 nL/min.

### Whole-mount clearing and imaging

The liver and trunk staining procedure was performed according to established protocols as previously described.^(14,48)^ In brief, the liver tissues were fixed in 4% paraformaldehyde (PFA), followed by permeabilization in PBS containing 0.2% Triton X-100, 20% DMSO, and 0.3 M glycine. Immunostaining was performed using the primary antibody against TH (Merck, Cat #AB152, 1:500), followed by incubation with a secondary antibody (Alexa Fluor 568-conjugated donkey anti-rabbit IgG, Thermo Fisher, Cat#A-10042). After staining, tissues were embedded in 1% agarose, dehydrated, and cleared using dibenzyl ether (DBE; Sigma, Cat#422053).

The brain-liver neurocircuit visualization was performed as previously described.^(25)^ At 5 days post-injection of PRV, the mice were anesthetized with isoflurane and perfused with PBS. Samples were permeabilized in PBS containing 1.5% goat serum, 0.5% Triton X-100, 0.5 mM methyl-β-cyclodextrin, and 0.2% trans-1-acetyl-4-hydroxy-l-proline. Following permeabilization, samples were stained with mRuby3 nanobody (NanoTag, Cat#N3302-AF647-L, 1:5,000). They were then dehydrated with upgraded tetrahydrofuran (THF; Sigma, Cat#401757), and subsequently incubated in dichloromethane (Sigma, Cat#270997). Finally, samples were cleared by benzyl alcohol and benzyl benzoate mixture (Sigma, Cats #24122 and #W213802).

The cleared samples were imaged using a light-sheet fluorescence microscope (Nuohai, LS18) at 1.25×, 3.2×, and 12.6× magnification. Finally, full-tissue image stacks were reconstructed and analyzed using Imaris ×64 software (version 9.5.0, Bitplane).

### Chemogenetic and optogenetic manipulations

For chemogenetic manipulation, we stereotaxically injected mice with 200 nL of AAV carrying the Cre-dependent activator DREADD (AAV9-DIO-hM3Dq-mCherry) and allowed them to recover for at least four weeks before conducting tests.

For optogenetic manipulation, mice received stereotaxic injections of 200 nL AAV9-DIO-ChR2-mCherry into the LPGi, followed by implantation of optical fibers (Inper, Cat#D3429). After a four-week recovery, mice were habituated to a fiber-optic patch cord in the home cage for 2 hours before the stimulation. Blue light (465 nm) stimulation, consisting of pulse trains (10 ms pulses of 20 Hz at approximately 10 mW; 1 s on, 1 s off), was delivered.

### Biochemical assays

The concentrations of epinephrine (EPI), norepinephrine (NE), insulin (INS) and glucagon (GC) were quantified using the EPI enzyme-linked immunosorbent assay (ELISA) kit (Kexing, Cat#Fankew F2351-A), NE ELISA kit (Kexing, Cat#Fankew F2533-A), INS ELISA kit (Kexing, Cat#Fankew F2579-A) and GC ELISA kit (Kexing, Cat#Fankew F2167-A). PEPCK activity assay kit (Kexing, Cat#Fankew F2533) and G6PC activity assay kit (Kexing, Cat#Fankew F2736-A) were used to quantify the activity of phosphoenolpyruvate carboxykinase (PEPCK) and glucose-6-phosphatase (G6PC).

### Western blot analysis

Tissue samples were homogenized in RIPA lysis buffer (Beyotime, Cat#P0013B) with 100 µM protease and phosphate inhibitors (Beyotime, Cat#P1045) using a homogenizer (JieChen Instrument, JCZYM-48R) at 4 °C. The following primary antibodies were used: PEPCK (Santa, Cat#sc-271029, 1:1,000), G6PC (Novus, Cat#NBP1-80533, 1:1,000), pPYGL (Ser15) (Abcam, Cat#ab227043, 1:1,000), PYGL (Proteintech, Cat#15851-1-AP, 1:1,000), pGS (Ser641) (Cell Signaling Technology, Cat#3891S, 1:1,000), GS (Santa, Cat#sc-81173, 1:1,000), TH (Merck, Cat#AB152, 1:1,000), and α-Tubulin (Huabio, Cat#ER130905, 1:10,000), followed by incubation with appropriate secondary antibodies. Imaging of membranes was performed with the iBright Imaging Systems (Thermo, iBright 1500). We quantified the signal intensities by performing densitometric analyses using Fiji (ImageJ, NIH).

### Histology and immunofluorescence analyses

Mouse livers and brains were collected separately and fixed with 4% PFA overnight at 4 °C. For Periodic Acid-Schiff (PAS) staining, 25 µm paraffin sections were prepared and stained with a PAS staining kit (Beyotime, Cat#C0142S). For immunostaining, 25-µm-thick frozen sections were prepared by a freezing microtome (Leica, CM 1950) and stained with primary antibodies, including anti-c-FOS (Synaptic Systems, Cat#226-008, 1:1,000), anti-TH (Merck, Cat#AB152, 1:500), anti-pPYGL (Ser15) (Abcam, Cat#ab227043, 1:100), and anti-pGS (Ser641) (Cell Signaling Technology, Cat#3891S, 1:100), followed by incubation with secondary antibody (Alexa Fluor 488-conjugated goat anti-rabbit IgG, Yeasen, Cat#33116ES60). The sections were imaged using confocal microscopes (Zeiss, LSM 700). Images were quantified with ImageJ (v1.8.0).

### Quantification and statistical analysis

An unpaired two-tailed Student’s t-test was used to compare differences between two groups. Comparisons across multiple groups were conducted using one-way ANOVA with the Tukey test. All results, including graphs, were expressed as means ± SEM unless otherwise specified. Significance was defined as *p* < 0.05, with significance annotations of \**p* < 0.05, \*\**p* < 0.01, \*\*\**p* < 0.001; ^#^*p* < 0.05, ^##^*p* < 0.01, ^###^*p* < 0.001; ^$^*p* < 0.05, ^$$^*p* < 0.01, ^$$$^*p* < 0.001. Statistical analyses were conducted using software such as GraphPad Prism (v.10.0), ImageJ (v1.8.0), and syglass (2.4.0).

## Abbreviations

AUC: area under the curve
CG: celiac ganglion
CGRP: Calcitonin gene-related peptide
CN: central nervous system
ChR2: channelrhodopsin-2
ELISA: enzyme-linked immunosorbent assay
EPI: epinephrine
G6PC: glucose-6-phosphatase
HPA: hypothalamic-pituitary-adrenal
LPGi: lateral paragigantocellular nucleus
NE: norepinephrine
6-OHDA: 6-hydroxydopamine
PAS: periodic acid-Schiff
PEPCK: phosphoenolpyruvate carboxykinase
PRV: pseudorabies virus
GS: glycogen synthase
PYGL: glycogen phosphorylase
WGA: wheat germ agglutinin.

## Data availability statement

All data generated or analyzed during this study are included in the manuscript and supporting files. Source data files have been provided for all figures.

## Author contribution

H.X., H.H., and X.Z. conceived, designed, and supervised the project. Z.W., M.R., Y.C., and Y.L. performed glucose level monitoring and data analysis. X.G., K.W., Y.C., and L.J. performed chemogenetic and optogenetic experiments. X.S., S.W., and J.W. performed whole-mount immunolabeling. H.X. and H.H. wrote the manuscript with input from H.W. and X.C. All authors validated and approved the final manuscript.

## Acknowledgement

We thank the following agencies for financial support: the National Natural Science Foundation of China (82301436, 82371311, 81930109, and 82321005), National Key R&D Program of China (2021YFA1301300, 2023YFA1801900); the Natural Science Foundation of Jiangsu Province (BK20221050, BG2024045), the Overseas Expertise Introduction Project for Discipline Innovation (G20582017001), the Jiangsu Specially Appointed Professor Program, and the International Postdoctoral Talent Recruitment Program, the Postgraduate Research & Practice Innovation Program of Jiangsu Province (KYCX25_1076). Graphical abstract was created with BioRender.com.

## Ethics approval and consent to participate

All animal experiments were approved by the Institutional Animal Care and Use Committee of China Pharmaceutical University and adhered to the principles outlined in the Guide for the Care and Use of Laboratory Animals.

