## Supplementary material for "Symmetric brain-liver circuits mediate lateralized regulation of hepatic glucose output in mice": Figure

### Figure S1

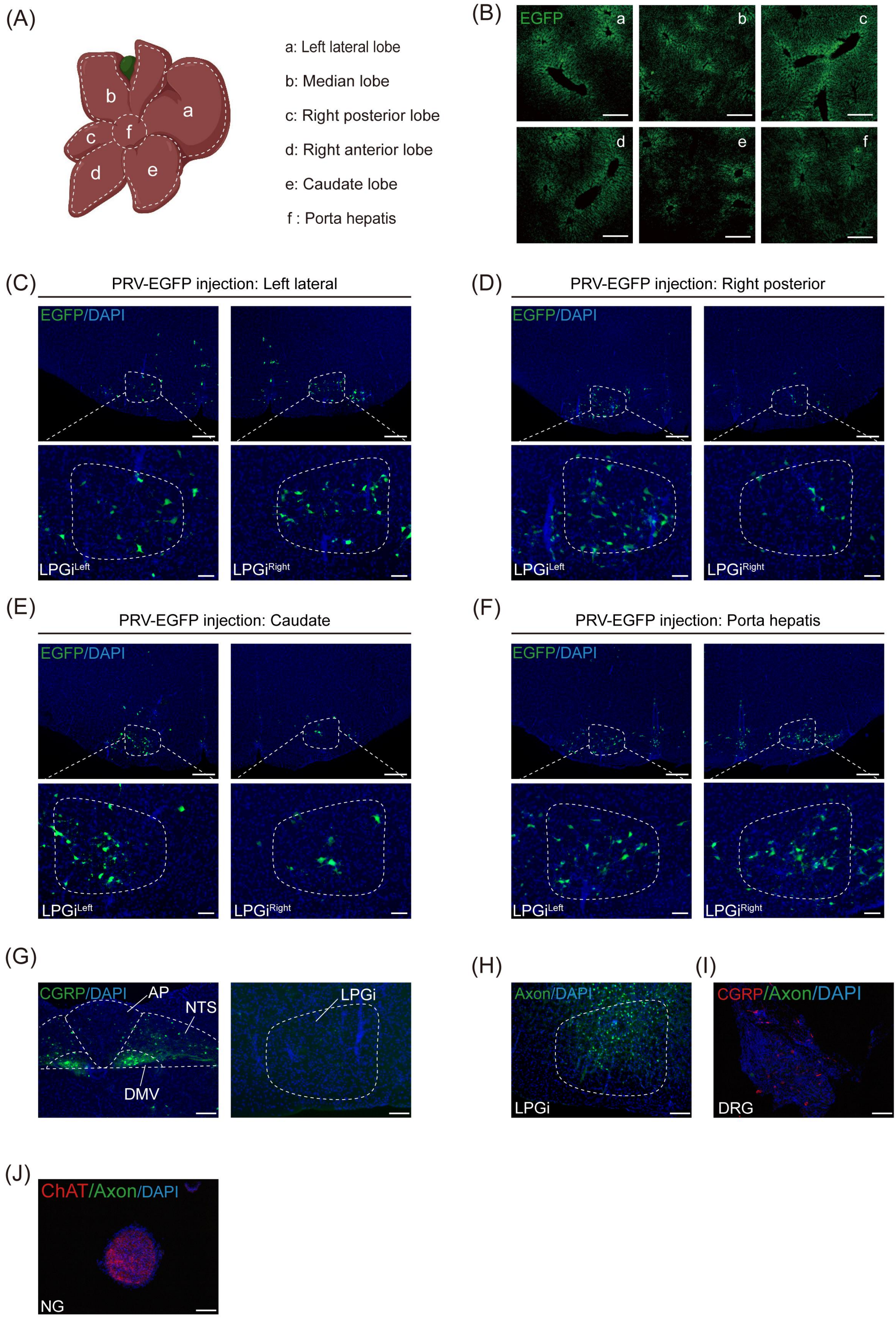
(A)
b
a
c
f
d
e
a: Left lateral lobe
b: Median lobe
c: Right posterior lobe
d: Right anterior lobe
e: Caudate lobe
f : Porta hepatis
(B)
EGFP
a
b
c
d
e
f
(C)
PRV-EGFP injection: Left lateral
EGFP/DAPI
LPGiLeft
LPGiRight
(D)
PRV-EGFP injection: Right posterior
EGFP/DAPI
LPGiLeft
LPGiRight
(E)
PRV-EGFP injection: Caudate
EGFP/DAPI
LPGiLeft
LPGiRight
(F)
PRV-EGFP injection: Porta hepatis
EGFP/DAPI
LPGiLeft
LPGiRight
(G)
CGRP/DAPI
AP
NTS
DMV
LPGi
(H)
Axon/DAPI
LPGi
(I)
CGRP/Axon/DAPI
DRG
(J)
ChAT/Axon/DAPI
NG

Supplementary figure 1. Supplementary evidence for contralateral projections from the brain to the liver, related to Figure 1.

(A) Schematic illustration of hepatic lobes with anatomical labeling.

(B) Representative images of hepatic lobes after PRV-EGFP injections. Scale bars, 150 μm.

(C-F) Representative immunofluorescence images of PRV-labeled neurons (EGFP) in the left and right LPGi following PRV injections into left lateral lobe (C), right posterior lobe (D), caudate lobe (E), and porta hepatis (F). Scale bars, 200 μm (top) and 100 μm (bottom).

(G) Representative immunofluorescence images showing CGRP (green) and DAPI (blue) in NTS (left) and LPGi (right). Scale bar , 100 μm

(H) Representative immunofluorescence images of Axon.-EGFP$^{+}$ neuron in LPGi. Scale bar , 100 μm

(I) Representative immunofluorescence images showing CGRP (red), Axon-EGFP labeling (green)and DAPI (blue) in DRG. Scale bar , 100 μm

(J) Representative immunofluorescence images showing choline acetyltransferase (ChAT, red),Axon-EGFP labeling (green) and DAPI (blue) inNG. Scale bar , 100 μm

### Figure S2

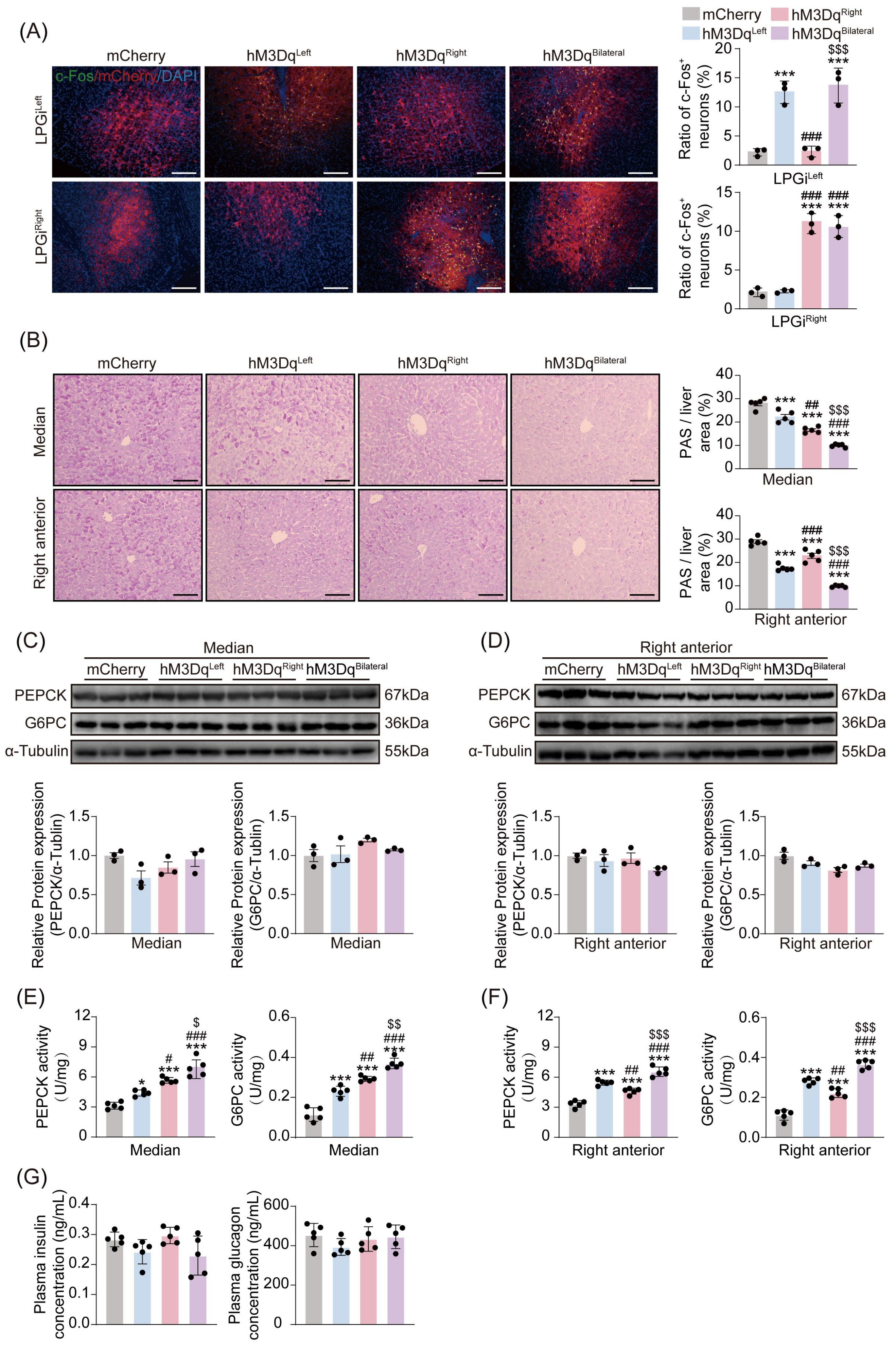
(A)
mCherry
hM3DqLeft
hM3DqRight
hM3DqBilateral
c-Fos/mCherry/DAPI
LPGiLeft
LPGiRight
Ratio of c-Fos+ neurons (%)
(B)
Median
Right anterior
PAS / liver area (%)
(C)
(D)
PEPCK
G6PC
α-Tubulin
67kDa
36kDa
55kDa
Relative Protein expression (PEPCK/α-Tublin)
Relative Protein expression (G6PC/α-Tublin)
(E)
(F)
PEPCK activity (U/mg)
G6PC activity (U/mg)
(G)
Plasma insulin concentration (ng/mL)
Plasma glucagon concentration (ng/mL)

Supplementary figure 2. Chemogenetic activation of LPGi induces c-Fos expression, promotes hepatic glycogenolysis, and activates glycogenolytic enzymes, related to Figure 2.

(A) Representative immunofluorescence images of c-FOS$^{+}$ LPGi neurons in hM3Dq mice following 1 h stimulation (Left), with quantification ofc-FOS$^{+}$ neuron percentage (right, $n$ = 3). Scale bars, 100 μm.

(B) Representative PAS staining images of median (top) and right anterior (bottom) lobes in hM3Dq mice following 1 h stimulation, with semi-quantitative analysis of glycogen levels (right, $n$ = 5). Scale bars, 100 μm.

(C and D) Western blot analysis of PEPCK and G6PC proteins in the median (C, top) and right anterior (D, top) lobes, with densitometric quantified (bottom, $n$ = 3).

(E and F) Enzyme activity of PEPCK and G6PC in the median (E) and right anterior (F) lobes ($n$ = 3).

(G) Plasma concentrations of insulin (Left, n = 5) and glucagon (Right, n = 5) $GAD^{mCherry}$ and $GAD^{hM3Dq}$ mice following 1h.

Data are expressed as means ± SEM, with individual values shown for all experiments. $^{*}p < 0.05$, $^{***}p < 0.001$; $^{\#\#}p < 0.01$, $^{\#\#\#}p < 0.001$; $^{\$\$\$}p < 0.001$. A one-way ANOVA with the Tukey test was used for (A-F).

### Figure S3

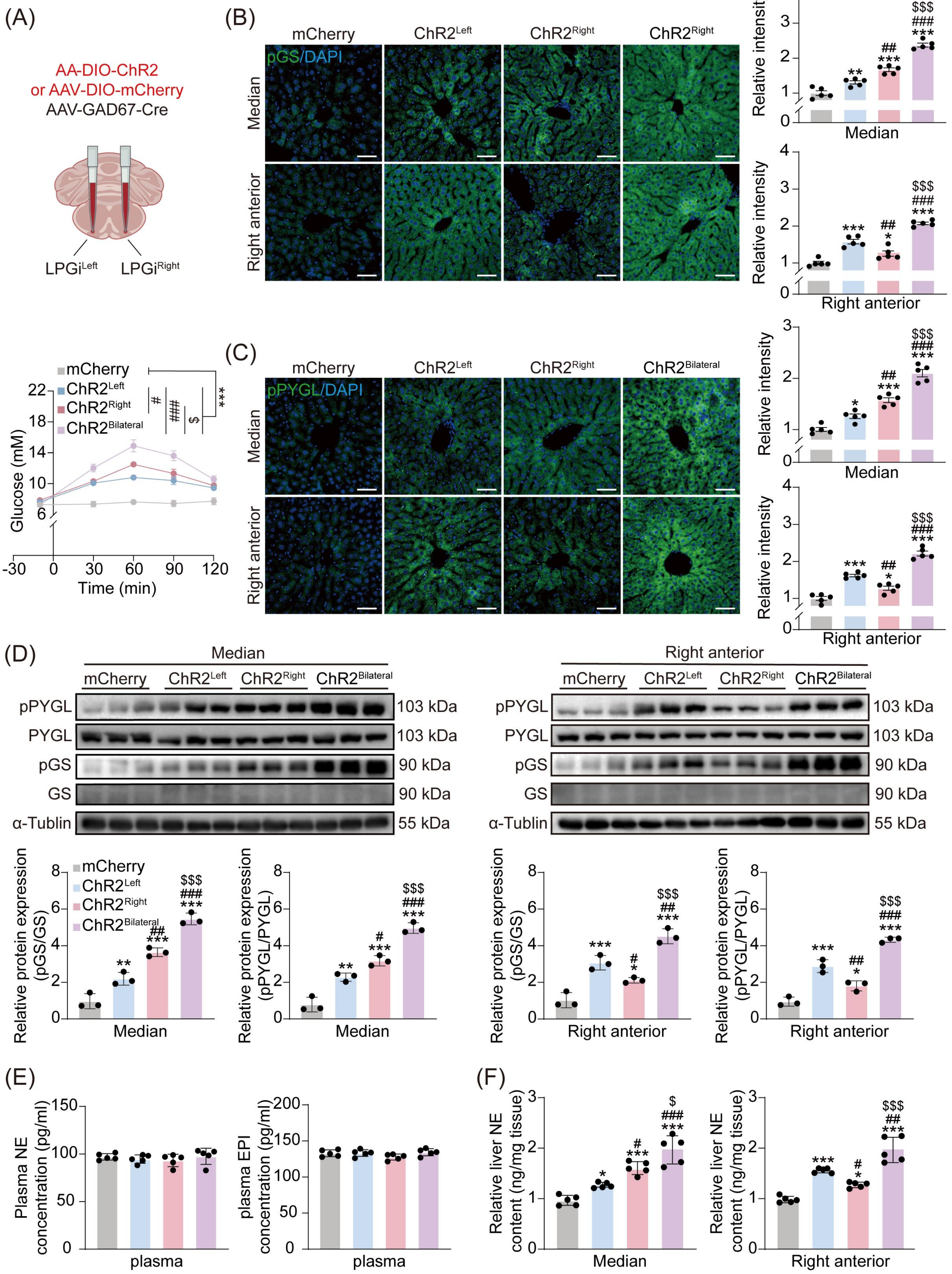

(A)
AA-DIO-ChR2
or AAV-DIO-mCherry
AAV-GAD67-Cre
LPGiLeft
LPGiRight
(B)
mCherry
ChR2Left
ChR2Right
ChR2Right
pGS/DAPI
Median
Right anterior
Relative intensity
Median
Right anterior
(C)
mCherry
ChR2Left
ChR2Right
ChR2Bilateral
pPYGL/DAPI
Median
Right anterior
mCherry
ChR2Left
ChR2Right
ChR2Bilateral
Glucose (mM)
Time (min)
(D)
Median
Right anterior
pPYGL 103 kDa
PYGL 103 kDa
pGS 90 kDa
GS 90 kDa
α-Tublin 55 kDa
Relative protein expression (pGS/GS)
Relative protein expression (pPYGL/PYGL)
(E)
Plasma NE concentration (pg/ml)
plasma EPI concentration (pg/ml)
plasma
(F)
Relative liver NE content (ng/mg tissue)
Median
Right anterior

Supplementary figure 3. Optogenetic activation of LPGi confirms contralateral lobe-specific mobilization observed in Figure 2.

(A) Scheme for injecting AAV-DIO-ChR2 or AAV-DIO-mCherry (top). Blood glucose levels following optogenetic activation of left, right, or bilateral LPGi, measured at −10, 30, 60, 90, and 120 min (bottom, $n = 6$)

(B and C) Representative immunofluorescence images of pGS (B) and pPYGL (C) expression in median (top) and right anterior (bottom) lobes after 1 h optogenetic activation of left, right, or bilateral LPGi, with quantification of relative fluorescence intensity ($n = 5$). Scale bars, 100 μm.

(D) Western blot analysis of pPYGL, PYGL, pGS, and GS in median (left) and right anterior (right) lobes after 1 h activation, with densitometric quantification (bottom, $n = 3$).

(E) Plasma concentrations of norepinephrine (NE) (left, $n = 5$) and epinephrine (EPI) (right, $n = 5$) following 1 h optogenetic activation.

(F) NE contents in median (left, $n = 5$) and right anterior (right, $n = 5$) lobes after 1 h optogenetic activation.

Data are expressed as means ± SEM, with individual values shown for all experiments. $^{*}p < 0.05$, $^{**}p < 0.01$, $^{***}p < 0.001$; $^{\#}p < 0.05$, $^{\#\#}p < 0.01$, $^{\#\#\#}p < 0.001$; $^{\$}p < 0.05$, $^{\$\$\$}p < 0.001$. A one-way ANOVA with the Tukey test was used for (A -F).

### Figure S4

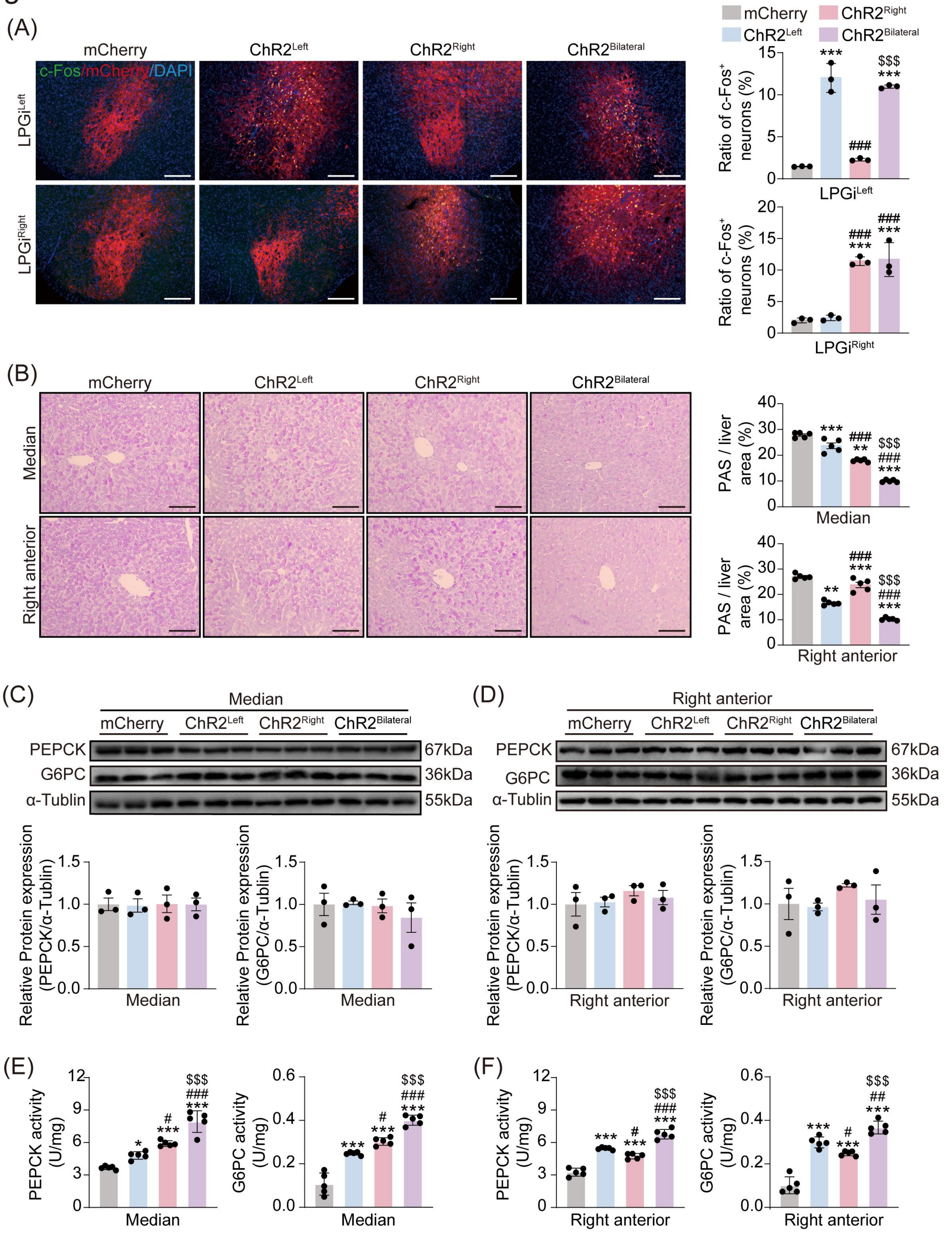

(A)
mCherry
ChR2Left
ChR2Right
ChR2Bilateral
c-Fos/mCherry/DAPI
LPGiLeft
LPGiRight
Ratio of c-Fos+ neurons (%)
(B)
Median
Right anterior
PAS / liver area (%)
(C)
(D)
PEPCK
G6PC
α-Tublin
67kDa
36kDa
55kDa
Relative Protein expression (PEPCK/α-Tublin)
Relative Protein expression (G6PC/α-Tublin)
(E)
(F)
PEPCK activity (U/mg)
G6PC activity (U/mg)

Supplementary figure 4. Optogenetic activation of LPGi recapitulates c-Fos expression, hepatic glycogenolysis, and activity of glycogenolysis-related enzymes observed in Figure S2.

(A) Representative immunofluorescence images of c-FOS$^{+}$ neurons in LPGi after 1 h optogenetic stimulation, with quantification of ratio of c-FOS$^{+}$ neurons (right, $n$ = 3). Scale bars, 100 μm.

(B) Representative PAS staining images of the median (top) and right anterior lobes (bottom) lobes following 1 h optogenetic stimulation, with semi-quantitative analysis of glycogen levels ($n$ = 5). Scale bars, 100 μm.

(C and D) Western blot analysis of PEPCK and G6PC proteins in median (C) and right anterior( D) lobes, with densitometric quantification ($n$ = 3).

(E and F) Enzyme activity of PEPCK and G6PC in median (E) and right anterior lobes (F) ($n$ = 3).

Data are expressed as means ± SEM, with individual values shown for all experiments. $^{*}p < 0.05$, $^{**}p < 0.01$, $^{***}p < 0.001$; $^{\#}p < 0.05$, $^{\#\#}p < 0.01$, $^{\#\#\#}p < 0.001$; $^{\$\$\$}p < 0.001$. A one-way ANOVA with the Tukey test was used for (A-F).

### Figure S5

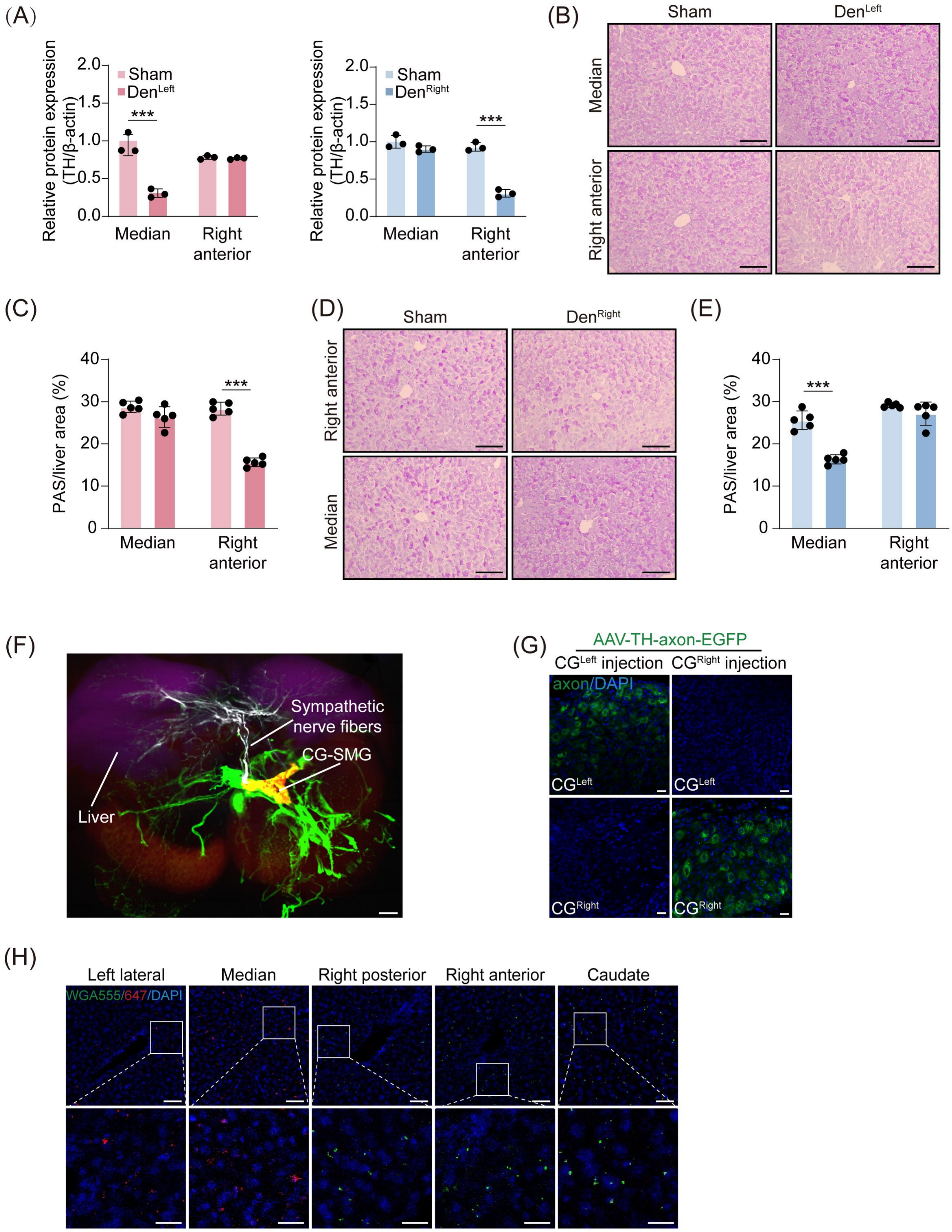

(A)
Relative protein expression (TH/β-actin)
2.0
1.5
1.0
0.5
0.0
Sham
DenLeft
***
Median
Right anterior
Sham
DenRight
***
(B)
Sham
DenLeft
Median
Right anterior
(C)
PAS/liver area (%)
40
30
20
10
0
***
Median
Right anterior
(D)
Sham
DenRight
Right anterior
Median
(E)
PAS/liver area (%)
***
Median
Right anterior
(F)
Sympathetic nerve fibers
CG-SMG
Liver
(G)
AAV-TH-axon-EGFP
CGLeft injection
CGRight injection
axon/DAPI
CGLeft
CGRight
(H)
Left lateral
Median
Right posterior
Right anterior
Caudate
WGA555/647/DAPI

Supplementary figure 5. Validation of sympathetic nerve ablation, enhanced glycogenolysis in non-denervated hepatic lobes, and accurate targeting of AAV and WGA injections, related to Figure 3 and 4.

(A) Densitometric quantification of TH protein in median and right anterior lobes after denervating left- (left) or right-sided (right) lobes.

(B-E) Representative PAS staining images of median (top) and right anterior (bottom) lobes in left- (B) and right-sided (D) lobes denervated mice, with semi-quantitative analysis of glycogen levels (C and E, $n = 5$). Scale bars, 100 μm.

(F) Representative 3D images of CG-SMG projections to peripheral organs (liver, purple; sympathetic nerve fibers, white; CG-SMG, yellow; TH-positive nerve fibers, green). Scale bar, 3,000 μm.

(G) Represent slices of the CG in unilateral AAV-TH-axon-EGFP injected mice. Scale bars, 20 μm.

(H) Represent slices of hepatic lobes after injecting WGA 647 and WGA 555 into left- and right-sided lobes. Scale bars, 50 μm.

Data are expressed as means ± SEM, with individual values shown for all experiments. $^{***}p < 0.001$; Unpaired Student's t-test for (A, C, and E).
